# DNA supercoiling impacts alternative transcription start site selection in yeast

**DOI:** 10.1101/2025.03.25.645220

**Authors:** Eleanor Elgood Hunt, Claudia Vivori, Folkert J van Werven

## Abstract

Most genes are transcribed from multiple transcription start sites (TSSs), defined as alternative TSSs, which are highly regulated and can lead to various gene regulatory outcomes including changes in translation efficiency and protein isoform expression. Transcription factors and chromatin regulators control alternative TSS selection. DNA supercoiling affects multiple aspects of transcription including transcription initiation; however, its regulatory effect on genes with multiple TSSs is not known. Here, we investigated how DNA supercoiling impacts alternative TSS usage in *Saccharomyces cerevisiae*. We depleted topoisomerases during early meiosis, where alternative TSS usage is prevalent, and applied an improved TSS sequencing protocol. We show that supercoiling affects alternative TSS usage at almost 600 genes. Increased alternative and aberrant TSS usage was observed near and within open reading frames, likely resulting from transcription-induced supercoiling originating from upstream alternative TSSs. DNA supercoiling had the greatest impact on genes with a dominant alternative TSS, significant spacing between alternative TSSs, and greater overall gene length. Our results establish that DNA supercoiling release during transcription is critical for correct TSS selection.

## Introduction

Most eukaryotic genes are transcribed from multiple transcription start sites (TSSs), defined as “alternative TSSs” (Carninci et al, 2006; FANTOM Consortium and the RIKEN PMI and CLST (DGT) et al, 2014). In budding yeast overlapping transcription units are common, with over 50% of genes expressing alternative TSSs (Pelechano et al, 2013; Lu & Lin, 2019). In humans, genes with alternative TSSs (estimated to be above 50%) are tightly regulated during development and are often misregulated in diseases such as cancer (Demircioğlu et al, 2019).

Variation in TSS usage is an impactful strategy to regulate gene expression. Alternative TSS usage can affect the 5’ untranslated region (5’ UTR), influencing translational efficiency, it can result in the production of distinct protein isoforms or in the synthesis of non-coding RNAs (Rojas-Duran & Gilbert, 2012; Fiszbein et al, 2019). A notable class of transcripts are long transcript undecodable isoforms (LUTIs): these transcripts originate from upstream promoters and are characterized by the presence of micro-upstream open reading frames (uORFs) in the 5′ leader sequence, which reduces their translation. In budding yeast, gene regulation by LUTIs is pervasive during meiosis, but also important for stress responses (Chen et al, 2017; Chia et al, 2017; Van Dalfsen et al, 2018; Tresenrider et al, 2021).

Alternative TSS usage is tightly regulated through mechanisms involving transcription factors and chromatin regulators (Chia et al, 2017; Tresenrider et al, 2021; Morse et al, 2024). Alternative TSSs can either be co-regulated or regulated by distinct transcription factors (Chia et al, 2021). Moreover, transcriptional interference from upstream TSSs can lead to the repression of downstream TSS usage and, as a consequence, lead to TSS switching events. The transcription inference by upstream transcription requires chromatin regulators. For example, the histone methyltransferase Set2, the chromatin remodelling complex FACT, and, more recently, the Swi/Snf2 complex have been shown to contribute to efficient switching between alternative TSSs, including of LUTIs (Chia et al, 2017; Tresenrider et al, 2021; Morse et al, 2024).

Transcription from one alternative TSS can influence the use of other TSSs, potentially through the effects of DNA supercoiling, which is known to impact transcription initiation. During transcription, positive supercoiling (or overwinding) and negative supercoiling (or underwinding) are generated in front and behind RNA polymerase (RNAP), respectively (Liu & Wang, 1987). Negative supercoiling can aid transcription by facilitating promoter melting, promoting transcription factor binding, and favouring the formation of the transcription complex (Parvin & Sharp, 1993; Tabuchi et al, 1993; Kouzine et al, 2013). Positive supercoiling can instead inhibit transcription initiation (Joshi et al, 2010). As transcription-produced supercoils can diffuse along DNA, transcription from one TSS may influence the transcription of nearby TSSs (Shearwin et al, 2005; Baranello et al, 2016; Asada et al, 2020; Patel et al, 2023). In tandem genes, transcription from an upstream TSS inhibits transcription at a downstream TSS due to positive supercoiling. Transcription from a downstream TSS instead promotes transcription of the upstream TSS due to negative supercoiling (El Hanafi & Bossi, 2000; Johnstone & Galloway, 2022).

Topoisomerases solve DNA topological problems by cutting the DNA, with type I topoisomerases cutting one DNA strand and type II topoisomerases cutting both strands (Baranello et al, 2014). In budding yeast, Top1 and Top2 act redundantly to enable RNAPII binding at transcribed genes, solving transcription-generated positive supercoiling and enabling efficient elongation (Sperling et al, 2011). Gene topology is regulated by Top2 at gene boundaries, while Top1 acts in coding regions (Achar et al, 2020). Co-depletion of Top1 and Top2 disrupts gene transcription predominantly at genes adjacent to one another.

While it is clear that DNA supercoiling affects multiple aspects of transcription, its role in regulating alternative TSS usage is unknown. We therefore hypothesised that transcription-induced supercoiling could affect alternative TSS selection. To investigate this, we co-depleted Top1 and Top2 in yeast cells staged in early meiosis, where alternative TSS usage is prevalent. Using an improved TSS sequencing method we showed that DNA supercoiling affected alternative TSS usage at almost 600 genes. DNA supercoiling had the largest effect on genes with a dominant alternative TSS and with large spacing between the alternative TSSs. Our results established that DNA supercoiling release during transcription is critical for correct alternative TSS selection.

## Results

### Efficiently induced depletion of Top1 and Top2 during early meiosis

Yeast meiosis provides an ideal system to study TSS regulation because alternative TSS usage is prevalent and its regulation is well characterized (Chia et al, 2021; Tresenrider et al, 2021). Moreover, cells can be synchronised to enter meiosis, which reduces confounding effects from asynchronous cell populations (Chia & van Werven, 2016). Previous works showed that Top1 and Top2 depletion can be used as an approach to induce genome-wide supercoiling (Achar et al, 2020; Patel et al, 2023). Therefore to determine how DNA supercoiling impacts alternative TSS usage, we generated Top1 and Top2 depletion alleles using the auxin-inducible degron (AID) system in cells (Figures 1A-1C) (Nishimura et al, 2009). We confirmed that the AID tag did not affect cell viability or the ability of cells to enter meiosis. Cells containing *TOP1-AID*, *TOP2-AID* and *TIR1* from the copper-inducible promoter showed comparable viability and meiotic onset to control cells (Figures 1D, S1A and S1B). When we co-depleted Top1 and Top2 (*TOP1/2-AID* and *TIR1*) by treatment with indole-3-acetic acid (IAA) and copper sulfate (CuSO_4_), we observed reduced cell viability and reduced number of cells that completed meiosis (71% vs 90% in control) (Figures 1D, S1A and S1B). The Top1+Top2 co-depletion strain (*TOP1/2-AID* + *TIR1*) is thus suitable for studying how DNA supercoiling affects alternative TSS usage in early meiosis.

**Figure 1.**
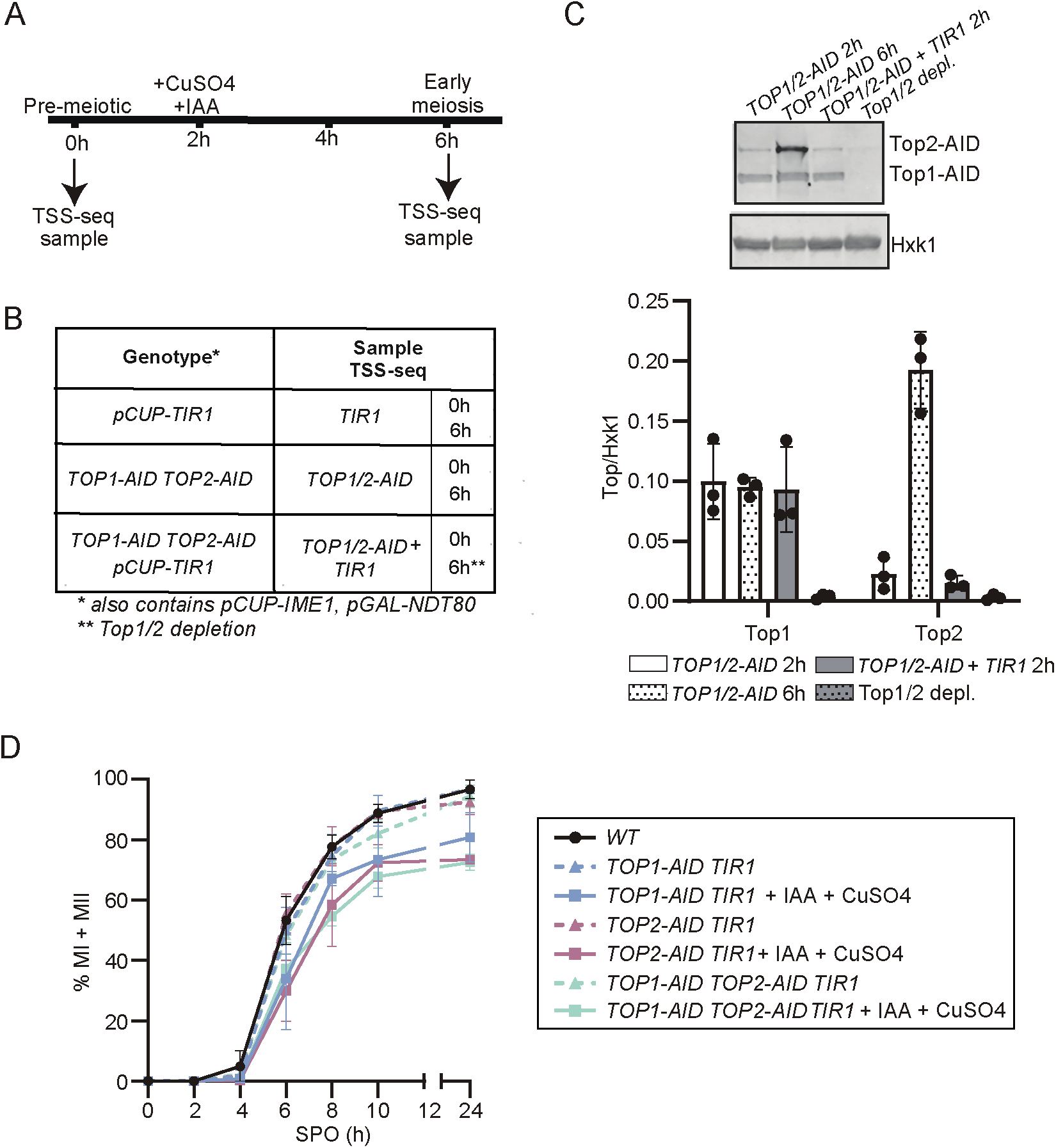
Co-depletion of Top1 and Top2 in early meiosis. **(A)** Schematic for experimental set-up. Cells were grown in rich media, then pre-sporulation medium, before being transferred to sporulation medium (SPO). Cells expressing pCUP-IME1, pGAL-NDT80 were used. Topoisomerase depletion and entry in meiosis were induced simultaneously by the addition of copper (II) sulphate (CuSO4) and indole-3-acetic acid (IAA) after 2 hours in SPO. NDT80 expression was not induced and so cells were arrested in early meiosis. TSS-sequencing samples were taken at 0 (pre-meiotic condition) and 6 hours (early meiosis) in SPO. **(B)** Table summarising genotypes, treatments and time points. TIR1 represents control strains only expressing pCUP-TIR1 (FW6109), TOP1/2-AID represents control strains only expressing AID-tagged Top1 and Top2 (FW11021), and TOP1/2-AID+TIR1 represents strains expressing both expressing pCUP-TIR1 and the AID-tagged Top1 and Top2 (FW11020). Before treatment, TOP1/2-AID+TIR1 cells still expressed Top1 and Top2, at 6 hours Top1 and Top2 were depleted (Top1/2 depl.), as indicated by asterisks. **(C)** Top1 and Top2 levels in the TOP1/2-AID control and TOP1/2-AID+TIR1 strains. Top1 and Top2 were detected using anti-V5 antibodies (the V5 epitope is fused to the AID tag), and the loading control Hxk1 was detected with anti-Hxk1 antibodies. Western blot quantification is shown by the bar plot (mean+SD, n=3 as shown by the black dots). **(D)** Effect of Top1/2 depletion on meiosis. Wild-type (WT) cells, cells expressing TOP1-AID TIR1, TOP2-AID TIR1, and TOP1-AID TOP2-AID TIR1 were used for the analysis (FW1511, FW10278, FW10279, FW12041). Cells were grown in rich media and subsequently in pre-sporulation medium, before being transferred to sporulation medium (SPO). and were either treated or not treated with IAA and CuSO4 at 2h SPO. Samples were taken at the indicated time points, fixed and stained with DAPI. Cells (n=200) with 2 or more DAPI masses were considered to have entered meiosis. Mean + SD of n = 3 experiments.

Next, we generated and tested Top1 and Top2 co-depletion alleles (*TOP1/2-AID* and *TIR1*) in cells staged in early meiosis (Figure 1A). High meiotic synchrony can be obtained by expressing the master regulatory transcription factor for meiotic entry, *IME1,* from the *CUP1* promoter (*CUP-IME1*) (Chia & van Werven, 2016). To ensure an arrest in early meiosis, we also introduced *NDT80* under the control of the *GAL1-10* promoter (*GAL-NDT80*). To induce meiosis, cells were grown to saturation in rich medium (YPD), overnight in pre-sporulation medium and then transferred to sporulation medium (SPO). After two hours in SPO, meiotic entry and Top1 and Top2 depletion were induced simultaneously by adding CuSO_4_ and IAA to the medium (Figure 1A). Since we did not induce *NDT80* expression, cells were arrested in early meiosis (6h SPO). Top1 and Top2 were co-depleted efficiently in early meiosis, achieving over 90% depletion (Top1/2 depl., Figure 1B and 1C). As controls, we used cells expressing either only TIR1 (*TIR*) or the *TOP1/2-AID* (*AID*) alleles (Figures 1B and 1C). Consistent with previous findings, Top2 expression increased during early meiosis, while Top1 expression was not changed in the *AID* control (Cheng et al, 2018). We conclude that Top1 and Top2 are efficiently co-depleted in early meiosis, which allows us to study the role of DNA supercoiling in alternative TSS usage regulation.

### Top1/2 depletion increases intragenic TSS usage

To accurately measure TSS usage, we applied an improved protocol for detecting and quantifying TSS usage at the single nucleotide resolution (TSS-seq) described in recently. In this protocol, we improved the reproducibility and quantification of TSS usage, optimising enzymatic reactions, and adapter sequences. Noteworthy, we used a 5’ adapter containing barcodes for sample multiplexing and unique molecular identifiers (UMIs). This allowed samples to be pooled after the adapter ligation step for further library preparation and enabled a quantitative assessment of TSS usage levels. The improved TSS-seq method was applied to samples at 0h SPO, representing the pre-meiotic stage, and 6h SPO, where all cells were staged in early meiosis.

We evaluated the accuracy of the TSS-seq technique by assessing the genome-wide locations of the identified TSSs. The detected TSS locations were compared to the annotated yeast open reading frames (ORFs). 77-79% of the identified TSSs at 0 hours were located in the promoter-proximal region (−300 +100 from annotated ORF start), while a lower proportion (68-70%) of the identified TSSs at 6 hours were located in this region, likely reflecting changes in TSS usage in meiosis (Figure 2A). As expected, the identified TSSs were predominantly located just upstream of the annotated yeast ORF start codons in the proximal promoter region (Figure S2A). The TSS-seq negative control (a sample not treated with the decapping enzyme) identified, as expected, fewer tags (more than 100-fold reduction) and these showed no enrichment for promoter-proximal regions (Figure S2B). Top1/2 depletion did not result in major global differences in the location of the identified TSSs, except for an increase in the proportion of identified TSSs located in the exonic regions (13%, compared to 10% in both controls) (Figure 2A).

**Figure 2.**
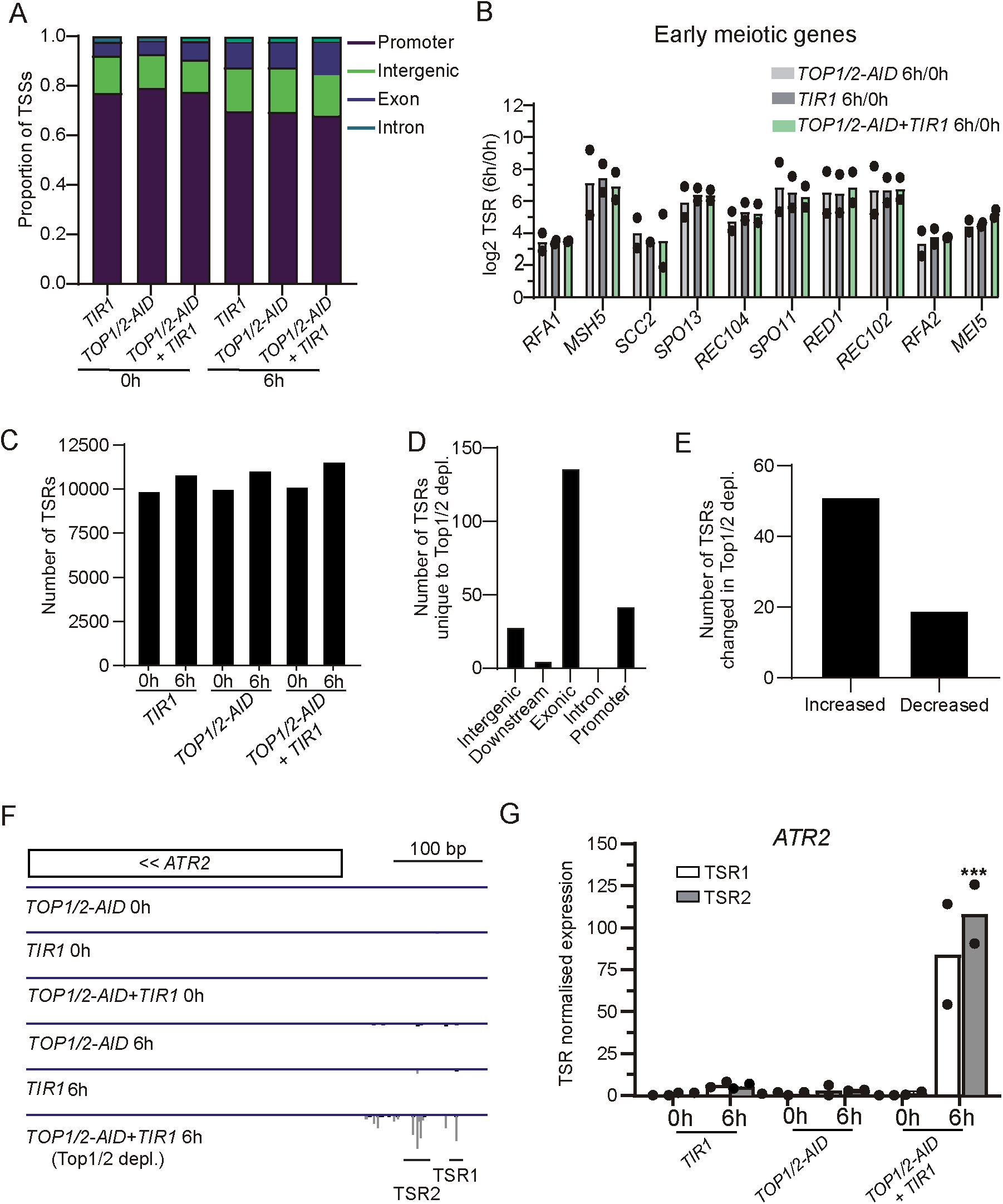
Top1/2 depletion affects TSR usage of a subset of transcripts. **(A)** Genomic locations of the detected TSSs at 0 and 6 hours in SPO for the different yeast strains. The promoter region was defined as −300 to + 100 bp from annotated yeast ORF start codons. n=2. **(B)** Bar plot showing the log2(fold-change) in the transcript start region (TSR) normalised expression levels of known early meiosis genes from 0 to 6 hours in sporulation media for the different strains. Tag counts (with a threshold of 3 counts) were normalised using power-law-based normalisation (Haberle et al, 2015) and tags within 5 bp were merged to identify TSRs. TSRs were annotated based on the closest gene on the same strand. For early meiotic genes with multiple annotated TSRs, the predominant TSR, within the promoter-proximal region, was plotted. Mean+SD, n=2 shown. **(C)** Bar plot showing the number of detected TSRs. TSRs less than 20 bp, and with a minimum normalised threshold of 1 were merged, and a set of non-overlapping consensus TSRs was generated. The number of TSRs with normalised read counts greater than 3 in both biological replicates of that sample was then quantified. **(D)** Bar plot showing the genomic locations of the TSRs only detected upon Top1/2 depletion (read counts greater than 3 in Top1/2 depl. 6h, and less than 3 in TIR1 6h and TOP1/2-AID 6h). **(E)** Bar plot summarising the number of TSRs usage at significantly different levels, at 6 hours, following Top1/2 depletion. The TSRs which significantly changed after Top1/2 depletion compared to both controls, but not between the control strains themselves, were counted. Calculated using DESeq2 with an FDR threshold of 0.05 and a fold-change threshold of 1.5. **(F)** Genome browser snapshot showing an example of TSS-sequencing profiles at 0 and 6 hours for the indicated strains at ATR2. Overlaid tracks from biological repeats for each strain are shown. **(G)** Bar plot showing the normalised ATR2 TSR1 and TSR2 usage (mean+SD, n=2 as shown by the black dots). Increased TSR usage was seen with Top1/2 depletion for ATR2: TSR1 increase was not significant (p=0.08), whilst TSR2 significantly increased (p=0.0001 as indicated by the asterisks). At 0 hours, similar TSR usage was seen between the strains.

Promoters initiate transcription at multiple positions, located close together, in TSS clusters or transcript start regions (TSRs) that are thus considered a part of the same core promoter (Batut et al, 2013). Following power-law-based normalisation, we grouped closely spaced TSSs (each located up to 5 bp apart) into TSRs, allowing comprehensive analysis of the TSS-seq data. The TSRs of known early meiotic genes (including *MSH5*, *SPO11* and *SPO13*), showed the expected increase in expression following meiotic onset, in both the control and Top1/2-depleted cells (Figure 2B) (Chia et al, 2021). Together, the data indicates that TSS-seq can accurately detect known TSSs and that meiosis was induced efficiently in the control and Top1/2 depletion strains.

To assess the correlation in TSR usage levels between samples, TSRs from all samples were aggregated (TSRs overlapping by less than 20 bp and with a minimum normalised read count of 1 were merged), to generate a set of non-overlapping consensus TSRs. At the pre-meiotic stage, where the samples should be equivalent, the detected TSRs showed a high correlation (Pearson’s r=0.89-0.97) (Figure S2C). The correlation between samples at 0 compared to 6 hours was lower, consistent with the expected changes in gene expression occurring during this cell fate transition. The correlation between the different strains was still high at 6 hours (Pearson’s r=0.95-0.98), suggesting that Top1/2 depletion did not cause global changes in TSR usage levels compared to the controls. Principal component analysis (PCA) also showed that the 0- and 6-hour samples clustered separately (Figure S2D).

The number of TSRs with a normalised read count of >3 in both biological replicates was further assessed (Figure 2C). Strikingly, more TSRs were detected in the Top1/2-depleted cells (11,566 compared to 10,823 for *TOP1/2*-*AID* 6h and 11,043 for *TIR1* 6h) (Figure 2C). The additional TSRs detected in the Top1/2-depleted cells were predominantly located inside ORFs (Figure 2D). This suggests that Top1/2 depletion increased intragenic transcription from TSSs.

To assess the effect of Top1/2 depletion, we performed a differential expression analysis of the consensus set of TSRs. A relatively small proportion of TSRs were significantly altered by Top1/2 depletion compared to the control strains (fold-change>1.5, FDR<0.05). 51 and 19 TSRs were significantly upregulated and downregulated upon Top1/2 depletion, respectively (Figures 2E, S3A-S3C). For example, at the *ATR2* locus, TSR1 and TSR2 usage increased by more than 100-fold upon Top1/2 depletion (Figure 2F and 2G).

### Top1/2 depletion affects alternative TSR usage at a subset of genes

Next, we assessed how Top1/2 depletion affected alternative TSR usage. First, we associated consensus TSRs with genes based on the nearest gene on the same strand. A gene was defined as having alternative TSRs if multiple TSRs were associated with the gene. We restricted the analysis to TSRs located within 1000 bp upstream of the start codon or within the coding region. Based on this definition, we found that 2,282 genes were associated with a single TSR, whilst 2,925 genes contained alternative TSRs (Figure 3A). Next, we calculated the alternative TSR ratio by determining the proportion of the most upstream TSR signal over the total signal from the other TSRs associated with the same gene, in the control and Top1/2 depleted samples. An increased ratio therefore meant more upstream TSR usage, while a decreased ratio indicated more downstream TSR usage (Figure 3B).

**Figure 3.**
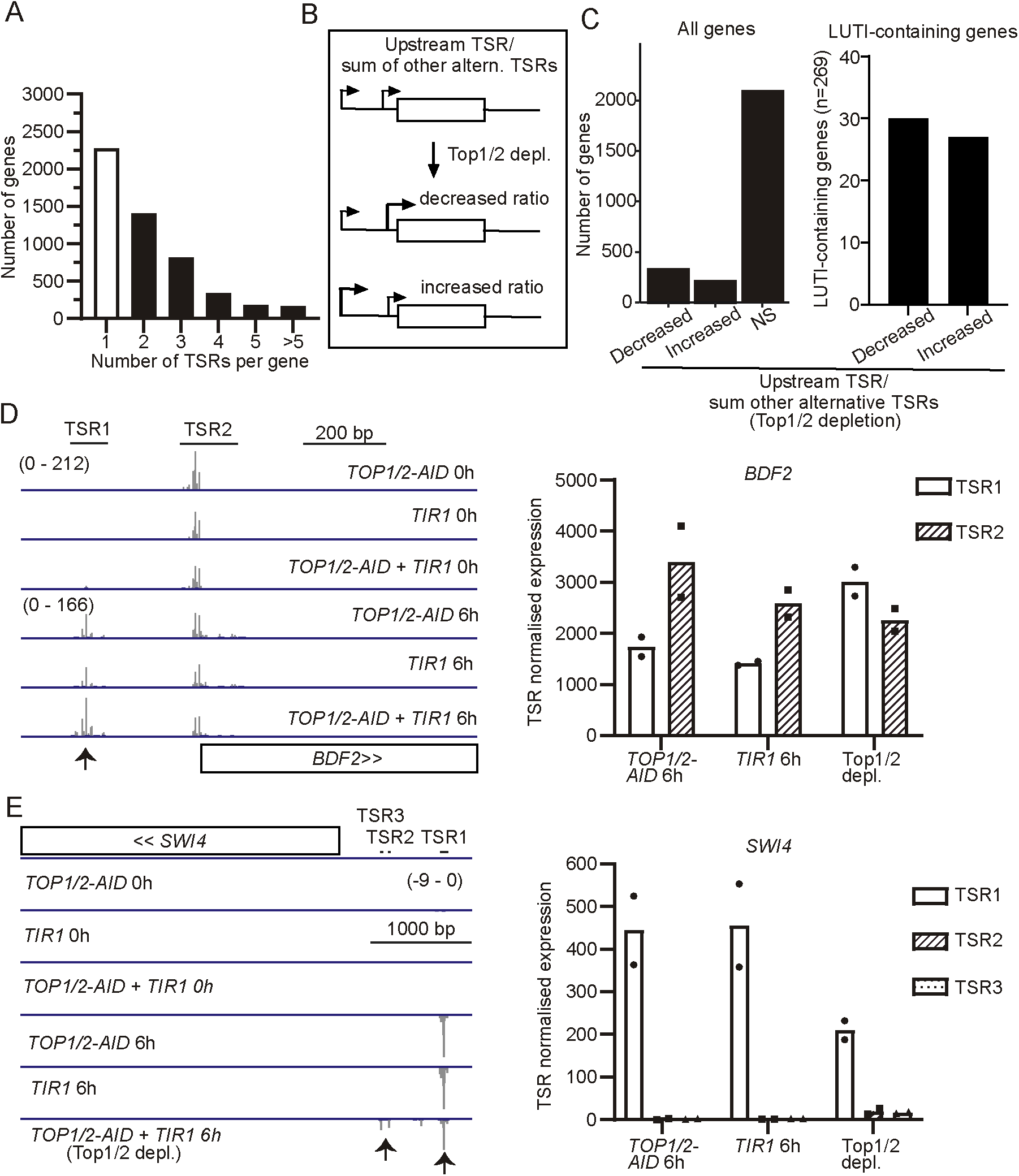
Top1/2 depletion affects alternative TSR usage. **(A)** Bar plot showing the number of genes associated with single or alternative TSRs. TSRs from all time points and samples were associated with genes based on the nearest gene on the same strand (with priority given to genes with ORF start codons within 1000 bp downstream of detected TSRs, and then genes with ORFs that overlapped the detected TSRs). A gene was defined as having alternative TSRs if multiple TSRs were associated with the gene (restricted to TSRs located up to 1000 bp upstream of the gene and including TSRs located internally to the gene). **(B)** Number of alternative TSR genes with significantly affected expression (p<0.001) of the 5’-most TSR compared to the total of all other TSRs, upon topoisomerase depletion. The genes with significant differences between the controls were removed from those significantly affected by topoisomerase depletion (NS; non-significant). Normalised read counts from DeSeq2 analysis were used, as done previously (Love et al, 2014; Chia et al, 2021). TSRs with normalised read counts above 1 in at least one 6-hour sample were filtered. The Cochran–Mantel–Haenszel test was used to assess statistical significance, across all biological replicates. **(C)** Bar plot showing the number of previously annotated long undecodable transcript isoforms (LUTI)-containing genes (Cheng et al, 2018) with relative alternative TSR expression ratio affected by topoisomerase depletion. 269 LUTI-containing genes were identified to express alternative TSRs by the TSS-sequencing technique. **(D)** Genome browser snapshot showing an example of TSS-sequencing profiles for BDF2 at 0 and 6 hours for the indicated strains. An overlay of the data from the biological repeats for each strain is shown. On topoisomerase depletion, the 5’-most TSR (TSR1) showed increased relative expression, compared to the control cells, as highlighted by the arrow. The normalised usage levels for each strain of BDF2 TSRs is shown by the bar plot. Mean+SD shown, n=2 (as shown by the black dots). **(E)** Genome browser snapshot showing an example of TSS-sequencing profiles for SWI4 (an example of a LUTI-expressing gene) at 0 and 6 hours for the indicated strains. An overlay of the data from the biological repeats for each strain is shown. On topoisomerase depletion, the 5’-most TSR (TSR1) showed decreased relative expression (as highlighted by the arrow). The normalised expression levels for each strain of SWI4 TSRs is shown by the bar plot. Mean+SD shown, n=2 (as shown by the black dots).

When comparing the *TIR1* control and the Top1/2-depleted cells, 283 genes showed an increased alternative TSR ratio, whilst 396 showed a decreased alternative TSR ratio (Cochran-Mantel-Haenszel test, Figure S4A). After correcting for both *TIR1* and *AID* controls, 237 and 354 genes showed increased and decreased alternative TSR ratios upon Top1/2 depletion, respectively (Figures 3C, left panel, and S4A). This gene set was used for further analyses.

A well-studied class of alternative TSSs are LUTI-containing genes (Chen et al, 2017; Chia et al, 2017; Cheng et al, 2018; Van Dalfsen et al, 2018; Tresenrider et al, 2021). LUTIs are characterized by transcription from upstream TSR leading to a long RNA isoform with an extended 5’UTR which is poorly translated due to uORFs in the leader sequence. During yeast meiosis, 380 genes were previously identified to express LUTIs (Cheng et al, 2018). We identified alternative TSRs at 269 of these LUTI-expressing genes. Upon Top1/2 depletion, 27 of the LUTI-expressing genes showed increased upstream alternative TSR usage, while 30 genes showed a decreased alternative TSR ratio (Figure 3C, right panel).

*BDF2* and *SWI4* are examples of alternative TSR-containing genes that are affected by Top1/2 depletion. *BDF2* showed an increased upstream TSR usage upon Top1/2 depletion (Figures 3D and S4B). *SWI4* is a LUTI-expressing gene with decreased upstream alternative TSR usage upon Top1/2 depletion (Figures 3E and S4C). We conclude that alternative TSR usage changes significantly at a large fraction of genes including a subset of LUTI-containing genes.

Next, we determined the location of the alternative TSRs affected by Top1/2 depletion. As expected, most upstream alternative TSRs were located in the promoter-proximal region (Figure S4D). For genes with a decreased alternative TSRs ratio upon Top1/2 depletion, the downstream TSRs were more frequently located within ORFs (Fisher’s exact test, p<0.0001, compared to genes with non-significant changes in alternative TSR usage) (Figure 4A and S4E). A similar result was obtained when we repeated the analysis for genes with two alternative TSRs only (Figure 4B). An example is the *MCM2* locus where depletion of Top1/2 leads to increased usage of four downstream TSRs, located in the ORF (Figures 4C, 4D and S4F). The analysis suggests that transcription-induced DNA supercoiling (induced by Top1/2 depletion) increases transcription from TSRs in gene ORFs.

**Figure 4.**
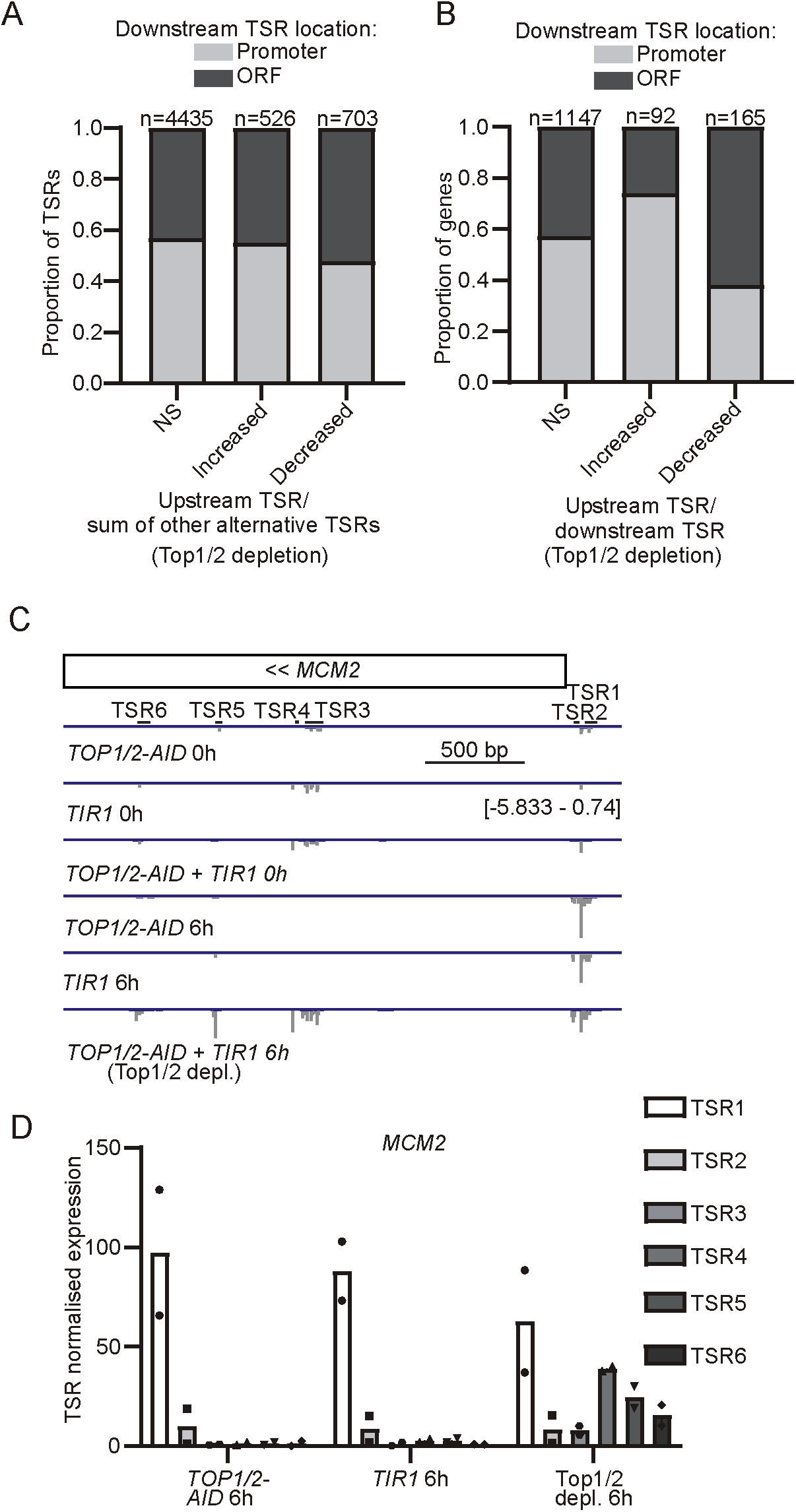
Top1/2 depletion affects relative alternative TSR usage of TSRs located in ORFs. **(A)** Bar plot showing the proportion of downstream TSRs located in the promoter-proximal region or ORF, for genes with relative alternative TSR expression ratios affected by topoisomerase depletion. The number of genes in each group is shown above the corresponding bar. **(B)** As in A, but with analysis restricted to genes with two alternative TSRs. Therefore, genes with increased relative upstream TSR usage will have decreased relative downstream TSR usage and vice versa. **(C)** Genome browser snapshot showing an example of TSS-sequencing profiles for MCM2 at 0 and 6 hours for the indicated strains. An overlay of the data from the biological repeats for each strain is shown. On topoisomerase depletion, the downstream TSRs, located in the exonic region of MCM2 were induced, and so decreased relative expression of the 5’-most TSR (TSR1) is seen, compared to the control cells. **(D)** Bar plot showing the normalised expression levels for each strain of MCM2 alternative TSRs. Mean+SD shown, n=2 (as shown by the black dots).

### Features of DNA supercoiling effects on alternative TSR usage

The varied outcomes of alternative TSR usage following Top1/2 depletion raised the question of whether specific regulatory features correlated with these differences. We reasoned that TSR usage levels and the distance between alternative TSRs or gene ends could impact the DNA supercoiling effects induced by Top1/2 depletion on TSR usage. We restricted our analysis to genes associated with two alternative TSRs to enable more direct comparisons between upstream and downstream TSR usage.

First, we examined whether a dominant downstream or upstream TSR, which likely generates stronger supercoiling, could exert more pronounced effects on upstream or downstream TSR usage respectively (Figure 5A). We found that genes with an increased TSR ratio upon Top1/2 depletion had a dominant downstream TSR (Wilcoxon rank sum test, 2-sided p<0.0001, compared to TSRs that did not significantly change) (Figure 5B). Conversely, genes with a decreased TSR ratio upon Top1/2 depletion had a dominant upstream TSR (Wilcoxon rank sum test, 2-sided p<0.0001, compared to TSRs that did not significantly change) (Figure 5B).

**Figure 5.**
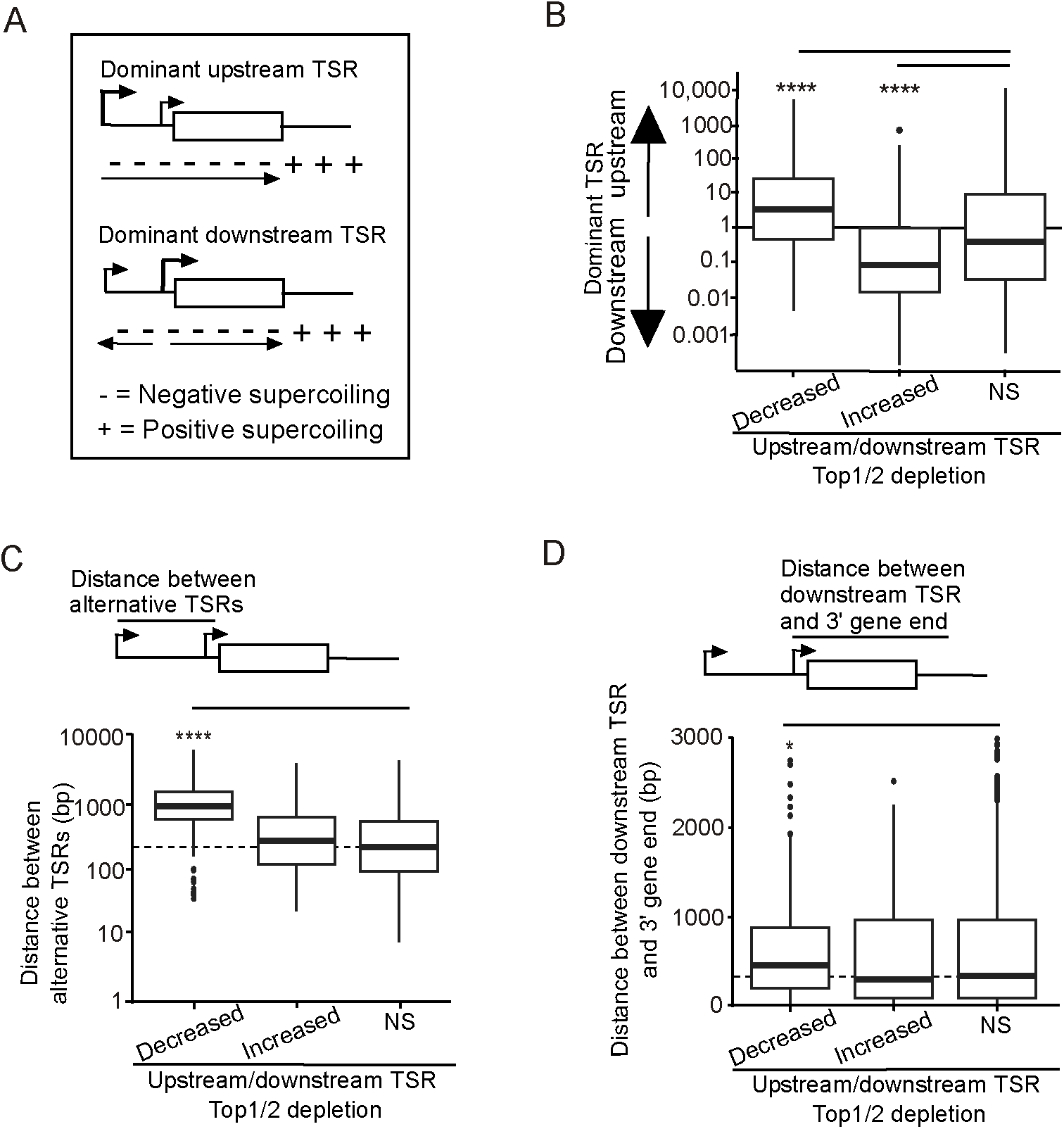
Regulatory features of alternative TSRs affected by Top1/2 depletion. **(A)** Model for how transcription-coupled DNA supercoiling affects alternative TSS usage. Transcription from upstream TSS may result in positive supercoiling in front of the RNAPII. When RNAPII has passed the downstream TSS, it is exposed to negative supercoiling. If the upstream TSS is dominant then the downstream TSS experiences stronger negative supercoiling. Negative supercoiling will also arise from downstream TSS transcription. If the downstream TSS is dominant, then the upstream TSS experiences stronger negative supercoiling. **(B)** Box plot showing the distribution of alternative TSR expression levels (in the control TIR cells at 6 hours) for genes with relative upstream TSR usage altered compared to downstream TSR usage upon topoisomerase depletion, with analysis restricted to genes with two alternative TSRs. Displayed box plots of genes with decreased (n=165), increased (n=92), or no significant changes (NS, n=1147) in relative alternative TSR usage (upstream over downstream TSR) after Top1/2 depletion. **(C)** Box plot showing the distribution of distances between alternative TSRs for genes with upstream TSR usage altered compared to downstream TSR usage upon Top1/2 depletion. Analysis was restricted to genes with two alternative TSRs. **(D)** Box plot showing the distribution of distances between downstream alternative TSR and 3’end of the gene. Analysis was restricted to genes with two alternative TSRs. Displayed box plots of genes with decreased, increased, or no significant changes in relative alternative TSR usage (upstream over downstream TSR) after Top1/2 depletion. Results of Wilcoxon rank sum test, compared to TSRs that did not significantly change, are indicated (* for P<0.05, ** for P<0.01, *** for P<0.001, **** for P<0.0001).

Second, we examined whether the distance between two TSRs affects TSR usage upon Top1/2 depletion. We reasoned that when the distance is greater, transcription from the upstream TSR possibly generates more pronounced supercoiling over the longer DNA distance, which may therefore have a stronger impact on downstream TSR usage. Indeed, genes with increased downstream TSR usage upon Top1/2 depletion displayed larger distances between the alternative TSRs (mean distance: 1000 bp versus 250 bp in the control, Wilcoxon rank sum test, 2-sided p<0.0001) (Figure 5C). Genes with an increased TSR ratio upon Top1/2 depletion instead showed no differences in the distance between alternative TSRs, compared to the control set.

Third, we examined whether gene length correlated with changes in TSR usage upon Top1/2 depletion. We computed the distance between the downstream TSR and 3’ gene end. We noted that for genes with increased downstream TSR usage upon Top1/2 depletion, the distance was significantly increased compared to the control (Wilcoxon rank sum test, 2-sided p<0.04) (Figure 5D). Genes with increased upstream TSR usage instead showed no significant difference in this distance compared to the control suggesting that upstream TSR usage was not affected by supercoiling effects at longer genes.

In conclusion, dominant TSRs, TSR spacing, and gene length correlate with changes to alternative TSR usage upon Top1/2 depletion. These data suggest that increased supercoiling can lead to complex regulatory outcomes in TSS usage.

## Discussion

Depletion of Top1 and Top2 is known to cause genome-wide induction of supercoiling at transcribed genes ^22^. Here we induced DNA supercoiling by co-depleting Top1 and Top2 in early yeast meiosis, where alternative TSS expression is prevalent, and showed that it impacts alternative TSS transcription at almost 600 genes, including LUTI-containing genes (Figure 3). Given the previously identified activatory effects of negative supercoiling, it is likely that transcription-induced negative supercoiling is responsible for the increased alternative TSS usage in ORFs. Our data further suggest that transcription-induced supercoiling mostly impacts alternative TSS usage at genes with dominant TSSs and with relatively large distances between alternative TSSs.

How does supercoiling regulate alternative TSS usage? In general, positive DNA supercoiling is thought to inhibit transcription and negative supercoiling to promote transcription (Pommier et al, 2016). However, there is also evidence that nucleosomes temper positive supercoiling and excessive negative supercoiling levels can inhibit transcription (Parvin & Sharp, 1993; Tabuchi et al, 1993; Kouzine et al, 2013; Herrero-Ruiz et al, 2021; Patel et al, 2023). Our data are consistent with transcription-induced negative supercoiling spreading over a distance to enhance alternative TSS usage. First, we find that Top1/2 depletion promotes alternative TSS usage at genes with a dominant alternative TSS (Figure 5B). Second, genes with larger distances between alternative TSSs were more affected by Top1/2 depletion. Specifically, we show that increased distances between the alternative TSRs increased downstream TSS usage (Figure 5C). During transcription from upstream to downstream TSSs, RNAPII spreads negative supercoiling when it passes the downstream TSS, which may open up chromatin and allow transcription factors access to activate downstream TSRs. Part of the mechanism could also include nucleosomes tempering initial excessive positive supercoiling levels when RNAPII is transcribing the region between TSSs, as has been shown by others (Teves & Henikoff, 2014; Corless & Gilbert, 2016).

Previous work, at tandem genes, showed that transcription from upstream TSSs generates positive supercoiling, which represses downstream TSS activity. Conversely, transcription from downstream TSSs induces negative supercoiling thereby enhancing transcription from upstream TSSs (Johnstone & Galloway, 2022). We show that alternative TSS-containing genes are partly regulated by a similar mechanism. Downstream alternative TSS usage induces negative supercoiling spreading to the upstream TSS. However, since RNAPII passes the downstream TSS, upstream TSS activity also promotes negative supercoiling at the downstream TSS (Figure 5).

Transcription-induced supercoiling may work in conjunction with histone modifications. Several modifications, including histone acetylation and methylation, have been linked to TSS usage changes (Chia et al, 2017; Moretto et al, 2018). Previously we showed that Set2-mediated histone H3 lysine 36 trimethylation (H3K36me3) and Set3-mediated histone deacetylation impact alternative TSS usage in meiosis, including at LUTI-containing genes (Chia et al, 2017; Tresenrider et al, 2021). We determined whether Top1/2 depletion affected alternative TSS usage at the same genes as single and double *set*2Δ *set*3Δ mutants. We noted little overlap between the two datasets (Figure S5). However, more work on dissecting the link between chromatin states, supercoiling events and alternative TSS usage is needed.

For this study, we did not measure the effect of depleting topoisomerases on supercoiling at genes with alternative TSSs. Various approaches have been used to study supercoiling events, including the use of psoralen, a reagent that intercalates in proportion to negative supercoiling levels (Achar et al, 2020; Yao et al, 2025). It may be insightful to apply some of these approaches; however, they typically do not capture the immediate states of transcription, which could present a challenge for the interpretation of the data.

Topoisomerases and DNA supercoiling play pervasive roles in gene transcription in mammalian cells, yet their impact on alternative TSS usage remains unclear. Notably, commonly used anticancer drugs function by inhibiting topoisomerase activity, and resistance to these drugs often arises through the downregulation of topoisomerase expression (Pommier et al, 2016). Understanding how topoisomerases regulate alternative TSS usage in mammalian cells could provide valuable insights into the potential side effects of these drugs.

## Materials and Methods

### Yeast strains and plasmids

All strains were derived from the Saccharomyces cerevisiae SK1 strain background. *TOP1* and *TOP2* were C-terminally tagged with the AID tag using a one-step tagging procedure as described previously (Longtine et al, 1998). pFA6a-3V5-IAA7-KanMx6 plasmid containing the 3xV5 and AID was used (Wu et al, 2018). The primers used for *TOP1* and *TOP2* tagging are shown in Table S2. Genetic crosses were employed to generate *TOP1-AID+TOP2-AID* together with *CUP1*-inducible *IME1* (*pCUP-IME1*) and *TIR1* (*pCUP-TIR*) as well as *GAL1-10* promoter *NDT80* strains as described in Figure 1 (Chia & van Werven, 2016). The genotypes of strains used are described in Table S1.

### Yeast growth conditions

Cells were arrested in early meiosis as described previously (Chia & van Werven, 2016). Briefly, yeast cells were inoculated in yeast extract peptone dextrose (YPD) media [1% weight/volume (w/v) yeast extract, 2% w/v bacto peptone, 2% w/v glucose, uracil (24 mg/l) and adenine (12 mg/l)] for approximately 24 hours, before being diluted to OD_600_ 0.4 in buffered, yeast extract, tryptone, acetate (BYTA, (1.0% w/v yeast extract, 2.0% w/v peptone, 1% potassium acetate, 50 mM potassium phthalate monobasic) and incubated for 16-18 hours. OD_600_ 1.8 of yeast cells were then washed and transferred to SPO (0.3% Potassium acetate, Acetic acid to pH 7.0, 0.02% D-(+)-Raffinose pentahydrate). 2 hours after transfer to SPO, CuSO_4_(50 μM) and IAA(500 μM) were added to induce synchronous meiotic entry and auxin-induced protein depletion. In all steps, yeast was grown at 30°C at 300 rpm, and a 1:10 culture volume to flask total volume ratio was maintained.

### Spot growth assay

Yeast cultures were grown in YPD for approximately 24 hours until saturation, before dilution to OD_600_ 0.4 in pure water. 5-fold serial dilutions were spotted onto rich growth media on agar plates with the addition of either 500 μM Dimethyl sulphoxide (DMSO) or 500 μM IAA and 50 μM CuSO_4_. Cells were imaged 2 days later.

### DAPI counting

At set time points following transfer of yeast cells to SPO, samples were collected by centrifugation, and fixed in 80% ethanol. Samples were then resuspended in 4’,6-diamidino-2-phenylindole (DAPI) (1 µg/ml in phosphate-buffered saline, PBS). The number of cells (n = 200) that had undergone one or two meiotic divisions (and so contained 2-4 nuclei) was assessed.

### Western blot

3.6 OD units of yeast were collected by centrifugation, fixed in 5% trichloroacetic acid (TCA), and incubated for a minimum of 10 minutes at 4°C. Cells were washed with acetone, and the pellet was dried. Cells were lysed using protein breaking buffer (50 mM Tris at pH 7.5, 1 mM EDTA, 27.5 mM DTT) and 0.5 mm glass beads (BioSpec) together with the Mini-Beadbeater-96 (BioSpec) for 5 minutes.

SDS loading buffer was added (62.5 mM Tris (pH 6.8), 2% β-mercaptoethanol, 10% glycerol, 3% SDS, and 0.017% Bromophenol Blue) and samples were boiled for 10 minutes at 100°C to denature the proteins. 4-20% gradient gels (Bio-Rad TGX) were used for SDS-PAGE (polyacrylamide gel electrophoresis), and protein was transferred to nitrocellulose membranes using Trans-Blot Turbo Transfer system (BioRad) as per the manufacturer’s instructions. Membranes were blocked with blocking buffer (1% w/v BSA, 1% w/v non-fat powdered milk in PBS with 0.01% v/v Tween-20 (PBST)) for at least 1 hour at room temperature, before incubation overnight at 4°C with primary antibodies (Table 3). Membranes were washed 3 times, 15 minutes, with 0.01% PBST, incubated with secondary antibodies in blocking buffer for 1 hour at room temperature, and then washed 3 times, 10 minutes.IRDye secondary antibodies were detected using an Odyssey Imager (Li-COR).

### RNA extraction

RNA was extracted from yeast as previously detailed (Chia et al, 2021). 24 OD units were collected by centrifugation, and snap-frozen in liquid nitrogen. Per 10 OD units, RNA was extracted with Tris-EDTA-SDS (TES) buffer (10 mM Tris-HCl pH 7.5, 10 mM EDTA, 0.5% SDS) and Acid Phenol:Chloroform:Isoamyl alcohol (125:24:1, Ambion) at 65°C for 45 minutes. After centrifugation at 4°C, max speed, 10 minutes, the aqueous phase was transferred to cold ethanol with 0.3 M sodium acetate. Precipitation was carried out overnight at 4°C. rDNase (Machery-Nagel) treatment was carried out for 20 minutes at 37°C, before spin column purification (Machery-Nagel).

### TSS-seq

The TSS-seq protocol used for this manuscript is described in a preprint. In short, total RNA was dephosphorylated with quickCIP (NEB, 1.2 U/µg RNA) 37°C, with RNasin Plus before heat inactivation. RNA was extracted using phenol/chloroform/isoamyl alcohol as before. RNA was treated with mRNA decapping enzyme (NEB, 0.7 U/µg RNA) at 37°C with RNasin Plus. RNA was extracted using phenol/chloroform/isoamyl alcohol. RNA and the 5’ adaptor (10 µM) were denatured for 2 minutes at 70°C, then cooled on ice for 2 minutes. Ligation was performed, 16°C using T4 RNA ligase I (30 U), with RNasin Plus. RNAClean XP (1.8X ratio, Beckman Coulter) was used to remove unligated adaptors, according to the manufacturer’s instructions. The concentration of the samples was measured using Qubit RNA BR assay, and samples were multiplexed. polyA+ RNA was enriched using Dynabeads Oligo(dT)_25_ (Thermo Fisher). Yeast RNA was fragmented with an alkaline fragmentation reagent (Ambion). RNeasy MinElute Clean-up columns, with 1.5x volume of Ethanol, were used to purify the samples. The 3’ ends were fixed by CIP treatment (NEB,30 U). Ligation was carried out by T4 Rnl2tr (NEB,200 U) for 2 hours, 25°C then 16 hours, 16°C. RNAClean XP (1.8X ratio) was used to remove unligated adaptors. RNA, RT oligo (0.5 pmol) and dNTP (10 mM each) were denatured at 65°C and then cooled. Reverse transcription was carried out by SuperScript III (2 µl), with RNasin Plus with the following conditions: 25°C for 5 minutes, 42°C for 20 minutes, 50°C for 40 minutes, 80°C for 5 minutes. Template RNA was removed by RNase H (10 U) at 37°C, for 30 minutes. AMPure XP beads (Agencourt, Beckman Coulter) were used to clean up the cDNA. 3X beads and 1.7x isopropanol were added to the reverse transcription reaction and incubated for 5 minutes. Beads were washed twice with 85% ethanol, dried, and eluted twice in nuclease-free water. cDNA pre-amplification was performed with i5_s and i7_s (300 nM each) primers and Phusion HF PCR Mastermix (Thermo Fisher). 6 PCR cycles were carried out (98°C for 10 seconds, 65°C for 30 seconds, 72°C for 30 seconds, with 3 minutes at 72°C final extension). ProNex beads (Promega) with a 1:2.95 ratio to sample were used to remove sequences less than 55 nt (including primer dimers). Final amplification was carried out with NEBNext i50 and i70 primers (500 nM each) and Phusion HF PCR Mastermix, for 8 cycles. Amplification was checked by using Novex 6% TBE (Tris/Borate/EDTA) gel and staining with SYBR green. ProNex beads (Promega) with a 1:2.3 ratio to sample were used for purification. Library concentration was measured using Qubit High Sensitivity D1000. Libraries were sequenced with Illumina NovaSeq 6000 using 100 nt paired-end reads, with 50 million reads minimum per sample.

### TSS-sequencing read mapping and trimming

Read processing was performed using an adapted nf-core/clipseq pipeline with Nextflow (Ewels et al, 2020). Read demultiplexing was performed using Ultraplex with parameters “--phredquality 15--min_length0” (Wilkins et al, 2021). Adaptor trimming was performed using Cutadapt with parameters “--minimum-length 20” (Martin, 2011). Bowtie2 was used for premapping to ribosomal and small RNAs (Langmead & Salzberg, 2012). Genome mapping was performed using STAR with the SK1-MvO-V1 annotation (Dobin et al, 2013). UMI-tools was used for deduplication (Smith et al, 2017). BedTools was used to identify the 5’-most nucleotide (tag) and for genome-wide comparative analyses (Quinlan & Hall, 2010).

### Downstream TSS-sequencing analysis

The majority of downstream analysis was performed using TSRexploreR and CAGEr (Haberle et al, 2015; Policastro et al, 2021). The promoter-proximal region was defined as −300 to +100 bp from annotated yeast start codons (based on SK1-MvO-V1 annotation for SK1).

For CAGEr analysis of yeast TSSs, 5’tag signals were normalised using Power-Law based normalisation (key parameters: α=1.12, T=10^6^)(Balwierz et al, 2009; Haberle et al, 2015). 5’ tags were clustered into TSRs (using clusterCTSS, with key parameters: method = “distclu”, maxDist = 5, keepSingletonsAbove = 3) (Chia et al, 2021). To assess correlation and perform differential expression, TSRs were aggregated to generate non-overlapping consensus TSRs (using aggregateTagClusters, with the key parameters: tpmThreshold = 1, qLow = 0.05, qUp = 0.95, maxDist = 20). Pearson’s correlation analysis between samples was performed using the normalised counts of each consensus TSR. DESeq2 was carried out on the consensus TSRs, using log2(fold-change)>=1, and FDR<0.05 (Love et al, 2014). TSRs were associated with the closest gene on the same strand. TSRs were annotated as exonic if they were located downstream of the annotated yeast ORF start codon. Relative enrichment for different gene classes was calculated, with Fisher’s exact test used to assess significance.

To assess the ratio of alternative TSR expression, consensus TSRs, from all time points and samples, were associated with genes based on the nearest gene on the same strand. Normalised read counts from DeSeq2 analysis were used, as done previously (Love et al, 2014; Chia et al, 2021). TSRs with normalised read counts above 1 in at least one 6-hour sample were selected. A gene was defined as having alternative TSRs if multiple TSRs were associated with the gene (restricted to TSRs located up to a maximum of 1000 bp upstream of the gene, and including TSRs located internally to the gene). For genes with alternative TSRs, the number of normalised reads associated with the most 5’ (upstream TSR) was divided by the number of reads associated with all other downstream TSRs. The Cochran–Mantel– Haenszel (CMH) test was used to assess whether this ratio was significantly different (p<0.001), across all biological replicates. The Wilcoxon rank sum test was used to assess whether genes with changed alternative TSR ratios were significantly associated with differences in relative expression level, inter-TSR distance, and gene length after the downstream TSR.

### Statistical analysis

Information regarding any statistical tests used, number of samples, or number of biological replicate experiments is stated in the corresponding figure legends. For t-tests, calculated P-values less than 0.05 were considered significant.

## Data Availability

The accession number for the RNA sequencing data reported in this paper is GSE292782.

## Acknowledgements

We thank the Crick Genomics STP for sequencing the libraries. We also thank the members of the van Werven lab for the critical reading of the manuscript. This work was supported by the Francis Crick Institute (CC2043), which receives its core funding from Cancer Research UK (CC2043), the UK Medical Research Council (CC2043), and the Wellcome Trust (CC2043).

