## Supplementary material for "DNA supercoiling impacts alternative transcription start site selection in yeast": Figures S1 - S5

Figure S1

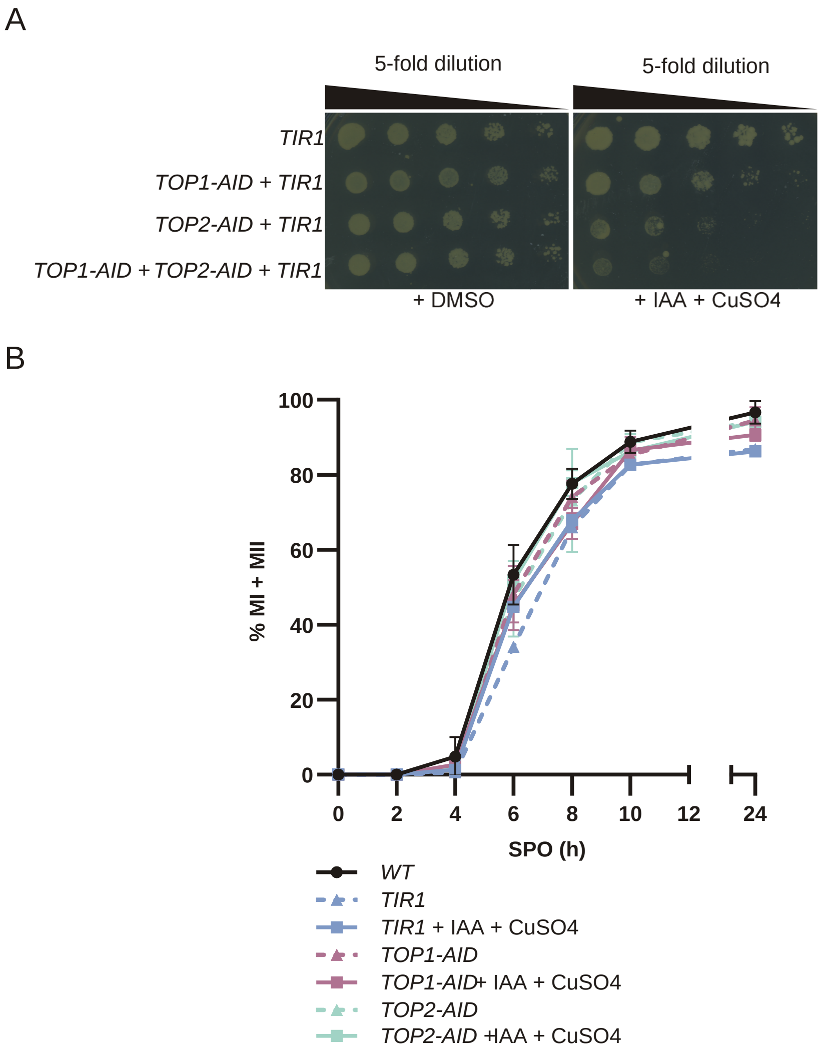


**Figure S1.** Co-depletion of Top1 and Top2 in early meiosis

**(A)** Spot growth assay to assess the effects on cell viability of the AID tag on Top1 and Top2 and Top1/2 induced depletion. 5-fold serial dilutions of strains were spotted onto rich growth media on agar plates with the addition of either DMSO or IAA and CuSO4. Strains expressing TIR1, TOP1-AID+TIR1, TOP1-AID+TIR2, and TOP1-AID+TOP1-AID+TIR1 were used (FW5737, FW10278, FW10279, FW12041). **(B)** Controls for the effect of Top1/2 depletion on meiosis. Wild-type (WT) cells, cells expressing TIR1, TOP1-AID or TOP2-AID were used for the analysis (FW1511, FW5737, FW12043, FW12044). Cells were grown in rich media and subsequently in pre-sporulation medium, before being transferred to SPO, and were either treated or not treated with IAA and CuSO4 at 2h in (SPO) Samples were taken at the indicated time points, fixed and stained with DAPI. Cells (n=200) with 2 or more DAPI masses were considered to have entered meiosis. Mean + SD of n = 3 experiments.

Figure S2

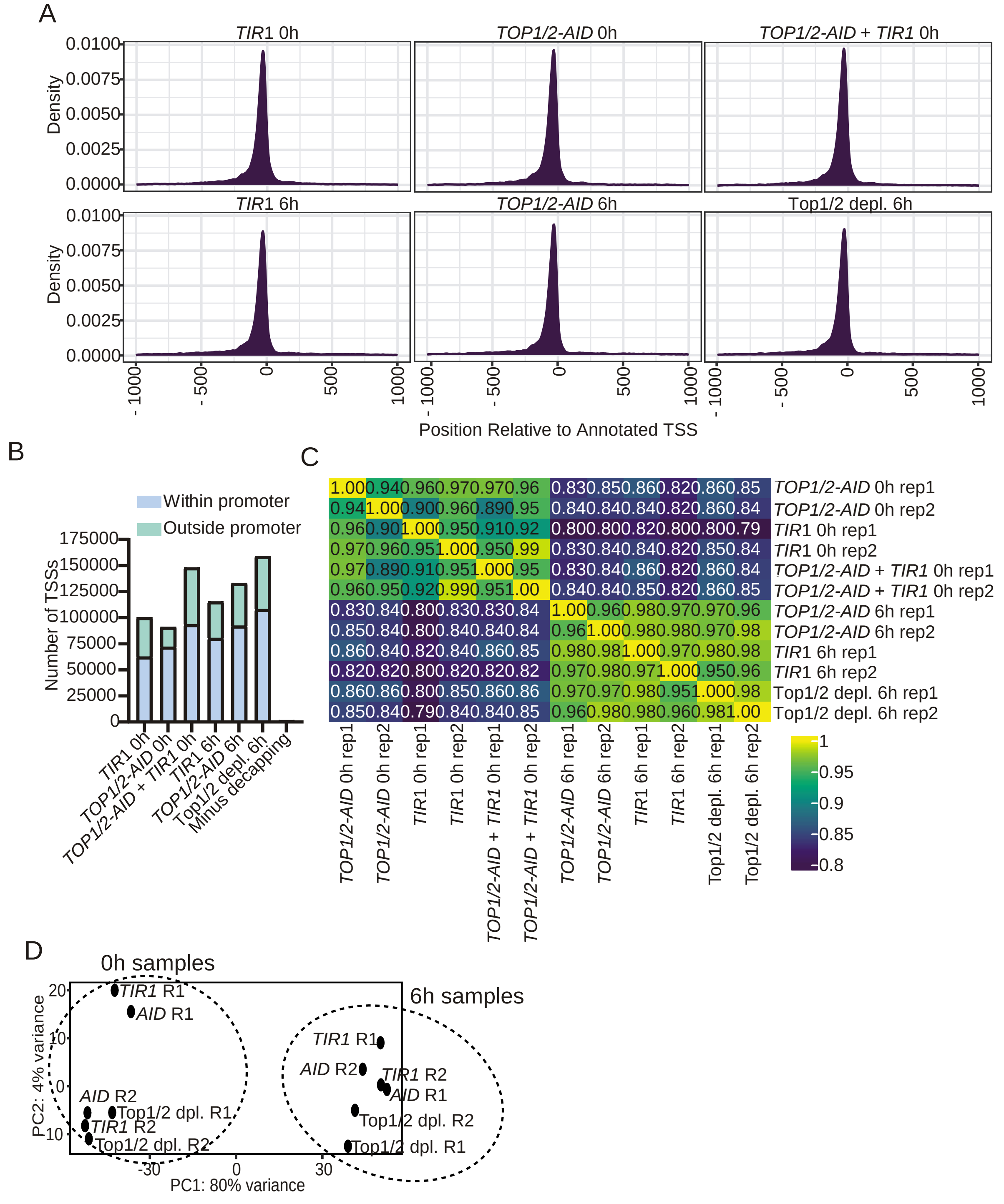


**Figure S2.** Clustering of TSS-sequencing data
**(A)** Density plots showing TSS locations compared to annotated yeast ORF start codons at 0 and 6 hours in sporulation media for the different yeast strains. **(B)** Bar plot showing the number of transcripts with promoter-proximal tags detected in the different samples. RNA that has not undergone the decapping enzyme reaction was used for the negative control as this RNA should be incompetent for adaptor ligation. **(C)** Heatmap of Pearson's correlation coefficients (r values) between the determined consensus TSRs of the different samples. **(D)** Principal component analysis of detected consensus TSRs for each sample. AID refers to TOP1/2-AID, and TOP refers to TOP1/2-AID+TIR1.

Figure S3

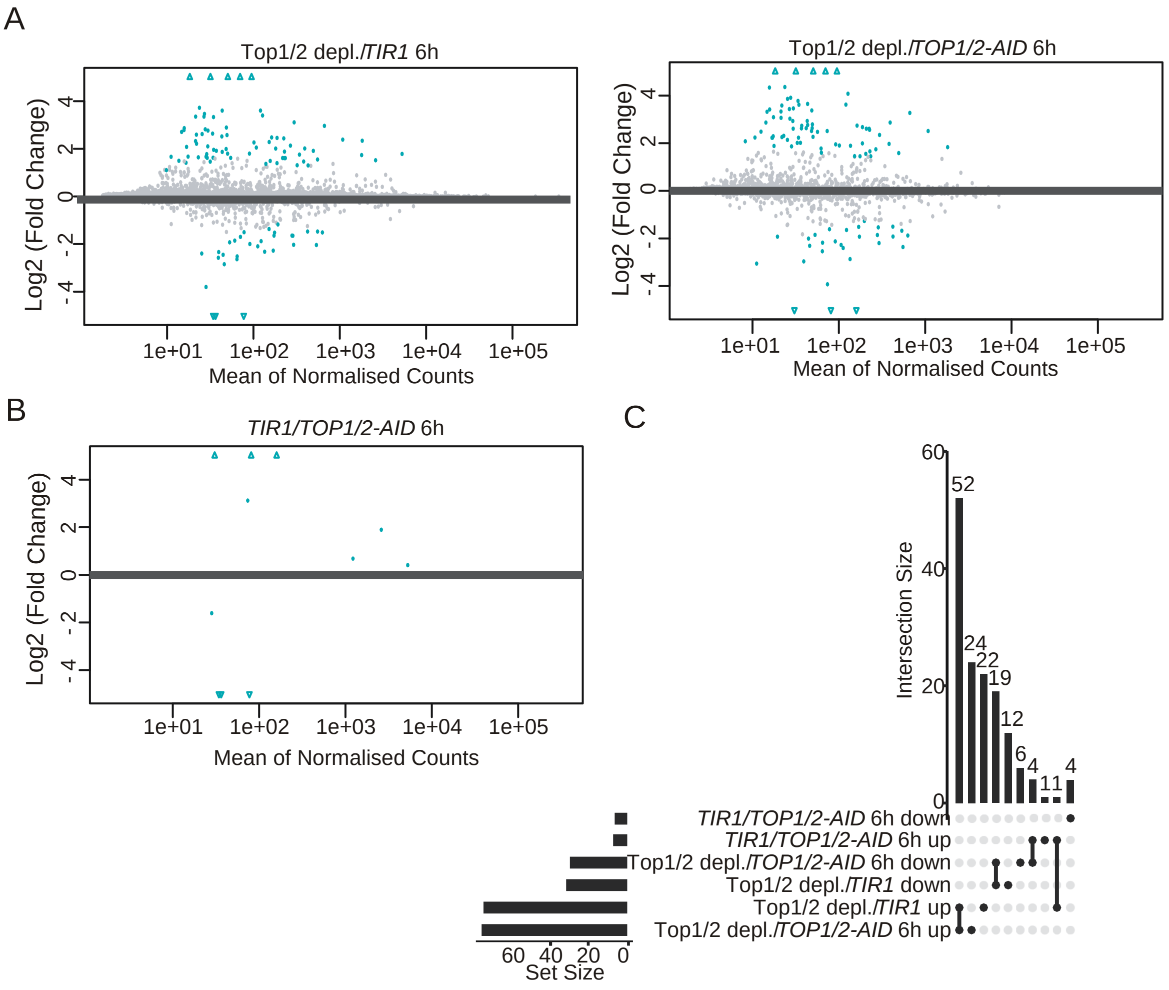


**Figure S3.** Effects of Top1/2 depletion on alternative TSR usage
**(A)** MA plots of differential consensus TSR usage at 6 hours between the Top1/2 depleted sample compared to the two control samples. Triangles indicate points outside of the axis limits. **(B)** MA plots of differential consensus TSR usage at 6 hours between the two control samples. Triangles indicate points outside of the axis limits. **(C)** Upset plot summarising the number of TSRs expressed at significantly different levels, at 6 hours, for the indicated comparisons. Calculated using DESeq2 with an FDR threshold of 0.05 and a fold-change threshold of 1.5.

Figure S4

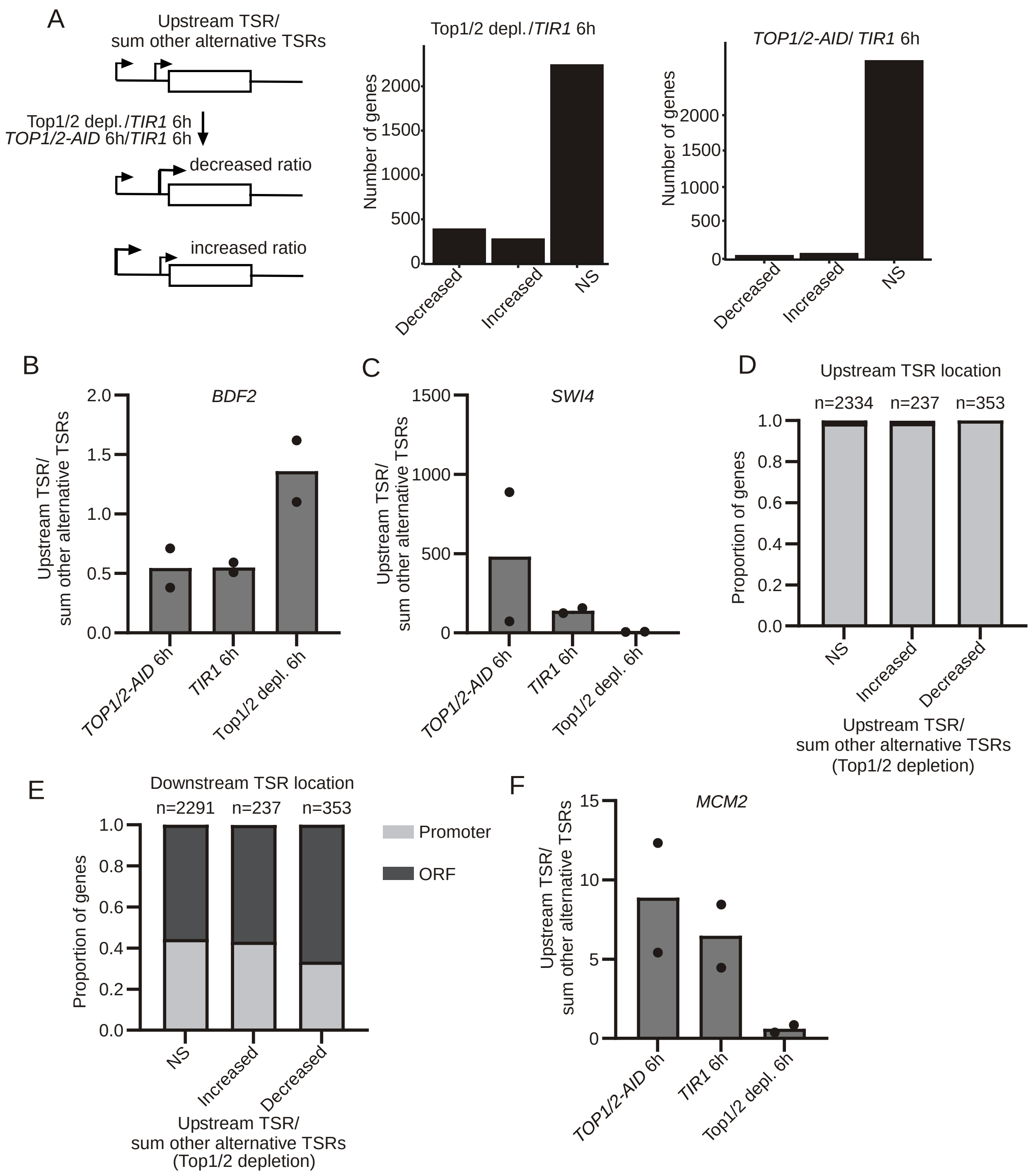


**Figure S4.** Effects of Top1/2 depletion on alternative TSR usage

**(A)** Number of alternative TSR genes with significantly affected expression (p<0.001) of the 5'-most TSR compared to the total of all other TSRs, in the indicated comparisons (NS, non-significant). Normalised read counts from DeSeq2 analysis were used. TSRs with normalised read counts above 1 in at least one 6-hour sample were filtered. A gene was defined as having alternative TSRs if multiple TSRs were associated with the gene (restricted to TSRs located up to a maximum of 1000 bp upstream of the gene, and including TSRs located internally to the gene). The Cochran–Mantel–Haenszel test was used to assess statistical significance, across all biological replicates. **(B)** Bar plot showing the ratio of expression levels of BDF2 TSR1 and TSR2. Mean+SD shown, n=2 (as shown by the black dots). **(C)** Bar plot showing the ratio of relative expression levels of SWI4 5'-most TSR and the sum of the other alternative TSRs. Mean+SD shown, n=2 (as shown by the black dots). **(D)** Bar plot showing the proportion of genes with 5'-most TSRs located in the promoter-proximal or ORF, for genes with relative alternative TSR expression ratios affected by topoisomerase depletion. The number of genes in each group is shown above the corresponding bar. **(E)** Bar plot showing the proportion of genes with all downstream TSRs located in the promoter-proximal region or at least one downstream TSR located in the ORF, for genes with relative alternative TSR expression ratios affected by Top1/2 depletion. The number of genes in each group is shown above the corresponding bar. (**F)** Bar plot showing the ratio of relative expression levels of MCM2 5'-most TSR and the sum of the other alternative TSRs. Mean+SD shown, n=2 (as shown by the black dots).

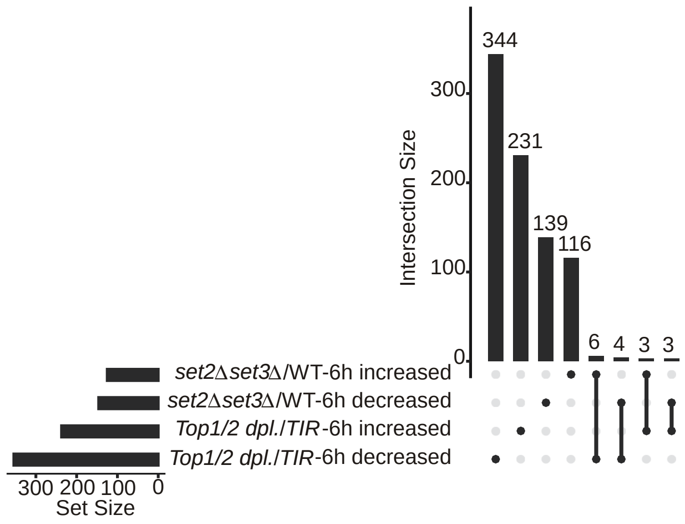


**Figure S5.** Set2 Set3 and Top1/2 control different alternative TSS-containing genes
Upset plot showing the number of genes with alternative TSR usage significantly affected by Set2 and Set3 depletion (Chia et al, 2021) and Top1/2 depletion in early meiosis (6h SPO).
