## Supplementary material for "DNA supercoiling impacts alternative transcription start site selection in yeast": Table S1

**Table S1.** Genotypes of the strains used.

| **Strain** | **Genotype** |
| --- | --- |
| *FW1509* | *MAT****a****, ho::LYS2, lys2, ura3, leu2::hisG, his3::hisG, trp1::hisG* |
| *FW1511* | *MAT****a****, ho::LYS2, lys2, ura3, leu2::hisG, his3::hisG, trp1::hisG*  *MATα, ho::LYS2, lys2, ura3, leu2::hisG, his3::hisG, trp1::hisG* |
| *FW2792* | *MAT****a****, ho::LYS2, lys2, ura3, leu2::hisG, his3::hisG, trp1::hisG*  *irt1::pCUP-3HA-IME1::HphMX , ndt80::pGAL-NDT80::TRP1, ura3::pGPD1-GAL4(848).ER::URA3* |
| *FW5377* | *MAT****a****, ho::LYS2, lys2, ura3, leu2::hisG, his3::hisG, trp1::hisG,*  *his3::pCUP-OsTIR:His3mx* |
| *FW5737* | *MAT****a****, ho::LYS2, lys2, ura3, leu2::hisG, his3::hisG, trp1::hisG,*  *his3::pCUP-OsTIR:His3mx*  *MATα, ho::LYS2, lys2, ura3, leu2::hisG, his3::hisG, trp1::hisG,*  *his3::pCUP-OsTIR:His3mx* |
| *FW6109* | *MAT****a****, ho::LYS2, lys2, ura3, leu2::hisG, his3::hisG, trp1::hisG, irt1::pCUP-3HA-IME1::HphMX , ndt80::pGAL-NDT80::TRP1, ura3::pGPD1-GAL4(848).ER::URA3, his3::pCUP-OsTIR:His3mx*  *MATα, ho::LYS2, lys2, ura3, leu2::hisG, his3::hisG, trp1::hisG, irt1::pCUP-3HA-IME1::HphMX , ndt80::pGAL-NDT80::TRP1, ura3::pGPD1- GAL4(848).ER::URA3, his3::pCUP-OsTIR:His3mx* |
| *FW10278* | *MATα, ho::LYS2, lys2, ura3, leu2::hisG, his3::hisG, trp1::hisG*  *Top1::Top1-3V5-IAA7::KanMX6. pNH603-pCUP-OsTIR*  *MAT****a****, ho::LYS2, lys2, ura3, leu2::hisG, his3::hisG, trp1::hisG*  *Top1::Top1-3V5-IAA7::KanMX6. his3:p550 (pNH603-pCUP-OsTIR)* |
| *FW10279* | *MATα, ho::LYS2, lys2, ura3, leu2::hisG, his3::hisG, trp1::hisG*  *Top2::Top2-3V5-IAA7::KanMX6. pNH603-pCUP-OsTIR*  *MAT****a****, ho::LYS2, lys2, ura3, leu2::hisG, his3::hisG, trp1::hisG*  *Top2::Top2-3V5-IAA7::KanMX6. his3:p550 (pNH603-pCUP-OsTIR)* |
| *FW10280* | *MAT****a****, ho::LYS2, lys2, ura3, leu2::hisG, his3::hisG, trp1::hisG*  *irt1::pCUP-3HA-IME1::HphMX , ndt80::pGAL-NDT80::TRP1, ura3::pGPD1-GAL4(848).ER::URA3, Top1::Top1-3V5-IAA7::KanMX6, his3::pCUP-OsTIR:His3mx* |
| *FW10281* | *MAT****a****, ho::LYS2, lys2, ura3, leu2::hisG, his3::hisG, trp1::hisG*  *irt1::pCUP-3HA-IME1::HphMX , ndt80::pGAL-NDT80::TRP1, ura3::pGPD1-GAL4(848).ER::URA3,Top2::Top2-3V5-IAA7::KanMX6, his3::pCUP-OsTIR:His3mx* |
| *FW11021* | *MAT****a****, ho::LYS2, lys2, ura3, leu2::hisG, his3::hisG, trp1::hisG, irt1::pCUP-3HA-IME1::HphMX , ndt80::pGAL-NDT80::TRP1, ura3::pGPD1-GAL4(848).ER::URA3, Top1::Top1-3V5-IAA7::KanMX6, Top2::Top2-3V5-IAA7::KanMX6*  *MATα, ho::LYS2, lys2, ura3, leu2::hisG, his3::hisG, trp1::hisG, irt1::pCUP-3HA-IME1::HphMX , ndt80::pGAL-NDT80::TRP1, Top1::Top1-3V5-IAA7::KanMX6, Top2::Top2-3V5-IAA7::KanMX6* |
| *FW11020* | *MAT****a****, ho::LYS2, lys2, ura3, leu2::hisG, his3::hisG, trp1::hisG, irt1::pCUP-3HA-IME1::HphMX , ndt80::pGAL-NDT80::TRP1, ura3::pGPD1-GAL4(848).ER::URA3, his3::pCUP-OsTIR:His3mx, Top1::Top1-3V5-IAA7::KanMX6, Top2::Top2-3V5-IAA7::KanMX6*  *MATα, ho::LYS2, lys2, ura3, leu2::hisG, his3::hisG, trp1::hisG, irt1::pCUP-3HA-IME1::HphMX , ndt80::pGAL-NDT80::TRP1, his3::pCUP-OsTIR:His3mx, Top1::Top1-3V5-IAA7::KanMX6, Top2::Top2-3V5-IAA7::KanMX6* |
| *FW12041* | *MAT****a****, his3D1, leu2D0, met15D0, ura3D0,Top1::Top1-3V5-IAA7::KanMX6, Top2::Top2-3V5-IAA7::KanMX6, his3::pGPD1-osTIR::HIS3*  *MATα, his3D1, leu2D0, met15D0, ura3D0,Top1::Top1-3V5-IAA7::KanMX6,Top2::Top2-3V5-IAA7::KanMX6,his3::pGPD1-osTIR::HIS3* |
| *FW12043* | *MATα, ho::LYS2, lys2, ura3, leu2::hisG, his3::hisG, trp1::hisG*  *Top2::Top2-3V5-IAA7::KanMX6.*  *MAT****a****, ho::LYS2, lys2, ura3, leu2::hisG, his3::hisG, trp1::hisG*  *Top2::Top2-3V5-IAA7::KanMX6.* |
| *FW12044* | *MATα, ho::LYS2, lys2, ura3, leu2::hisG, his3::hisG, trp1::hisG*  *Top1::Top1-3V5-IAA7::KanMX6.*  *MAT****a****, ho::LYS2, lys2, ura3, leu2::hisG, his3::hisG, trp1::hisG*  *Top1::Top1-3V5-IAA7::KanMX6.* |
