## Supplementary material for "DNA supercoiling impacts alternative transcription start site selection in yeast": Table S2

Table S2. Sequences of oligos primers used.

| *TOP1*-Fw | 5’-CAAATGGGCCATAGAATCGGTAGATGAAAATTGGAGGTTTcggatccccgggttaattaa |
| --- | --- |
| *TOP1*-Rv | 5’-TGAATGTATTTGCTTCTCCCCTATGCTGCGTTTCTTTGCGgaattcgagctcgtttaaac |
| *TOP2*-Fw | 5’-GGAAAACCAAGGATCAGATGTTTCGTTCAATGAAGAGGATcggatccccgggttaattaa |
| *TOP2*-Rv | 5’- ACATATAAAAAGAATGGCGCTTTCTCTGGATAAATATTATgaattcgagctcgtttaaac |
