## Supplementary material for "DNA supercoiling impacts alternative transcription start site selection in yeast": Table S3

**Table S3. Antibodies used for Western blot analysis**

| **Antigen** | **Host** | **Source** | **Catalog#** | **Dilution** |
| --- | --- | --- | --- | --- |
| V5 | Mouse (monoclonal) | Invitrogen | AB_2556564 | 1:2000 |
| Hexokinase | Rabbit (polyclonal) | Stratech | H2035 | 1:8000 |
| Anti-mouse IgG, HRP-linked | Sheep | GE Life Sciences | NA931V5 | 1:10000 |
| Anti-rabbit IgG, HRP-linked | Donkey | GE Life Sciences | NA934V | 1:10000 |
| IRDye 800CW anti-mouse | Goat | LiCOR | 926-32210 | 1:15,000 |
| IRDye 680RD anti-Rabbit | Goat | LiCOR | 926-68071 | 1:15,000 |
